# A bacterial symbiont protects honey bees from fungal disease

**DOI:** 10.1101/2020.01.21.914325

**Authors:** Delaney L. Miller, Eric A. Smith, Irene L. G. Newton

**Affiliations:** Department of Biology, Indiana University, Bloomington, Indiana, USA

## Abstract

Fungi are the leading cause of insect disease, contributing to the decline of wild and managed populations^1,2^. For ecologically and economically critical species, such as the European honey bee (*Apis mellifera*), the presence and prevalence of fungal pathogens can have far reaching consequences, endangering other species and threatening food security^3,4,5^. Our ability to address fungal epidemics and opportunistic infections is currently hampered by the limited number of antifungal therapies^6,7^. Novel antifungal treatments are frequently of bacterial origin and produced by defensive symbionts (bacteria that associate with an animal/plant host and protect against natural enemies ^89^. Here we examined the capacity of a honey bee-associated bacterium, *Bombella apis*, to suppress the growth of fungal pathogens and ultimately protect bee brood (larvae and pupae) from infection. Our results showed that strains of *B. apis* inhibit the growth of two insect fungal pathogens, *Beauveria bassiana* and *Aspergillus flavus, in vitro*. This phenotype was recapitulated *in vivo*; bee brood supplemented with *B. apis* were significantly less likely to be infected by *A. flavus*. Additionally, the presence of *B. apis* reduced sporulation of *A. flavus* in the few bees that were infected. Analyses of biosynthetic gene clusters across *B. apis* strains suggest antifungal production via a Type I polyketide synthase. Secreted metabolites from *B. apis* alone were sufficient to suppress fungal growth, supporting this hypothesis. Together, these data suggest that *B. apis* protects bee brood from fungal infection by the secretion of an antifungal metabolite. On the basis of this discovery, new antifungal treatments could be developed to mitigate honey bee colony losses, and, in the future, could address fungal epidemics in other species.

---

Emerging fungal pathogens pose major threats to animal and plant populations^2^. Among insects, fungal pathogens are currently the most common causal agents of disease, and historically have plagued insect hosts for over 300 million years^1,10^. In recent years, fungal pathogens have contributed to the unprecedented population decline of honey bees, causing opportunistic infections in already stressed colonies ^3,4^. Within the colony, the most susceptible individuals are arguably the bee brood (larvae and pupae), which are exposed to fungal pathogens, notably chalkbrood (*Ascophaera apis*) and stonebrood (*Aspergillus flavus*) ^11,12^. Although the spread of fungal disease among the brood can be limited by the hygienic behavior of honey bee nurses^13^, this behavior does not prevent infection. However, brood fungal infections in other insects are sometimes inhibited by the presence of bacterial symbionts^14,15,8^. Given that honey bee brood are reared in the presence of a handful of bacterial taxa^16,17^, it is possible these microbes play similar defensive roles. Indeed, worker honey bee pathogen susceptibility correlates with changes in their microbiome composition and abundance ^18,19,20,21^. Furthermore, the presence of key microbiome members in worker bees can alter the prevalence of bacterial diseases ^22,23,24,25^. In aggregate, this evidence suggests that honey bee-associated bacteria can defend against bacterial pathogens and may similarly protect the host from fungal disease.

One of the most prevalent brood-associated bacteria is *Bombella apis* (formerly *Parasaccharibacter apium*), an acetic-acid bacterium found in association with nectar and royal jelly. Within the colony it is distributed across niches including larvae, the queen’s gut, worker hypopharyngeal glands, and nectar stores. Many of the niches it colonizes, particularly the larvae, are susceptible to fungal infection and/or contamination, and its localization to these niches may be indicative of a protective role. Furthermore, increased *B. apis* load is negatively correlated with *Nosema* (a fungal pathogen) in honey bee adults, suggesting interactive effects. However, since *B. apis* is rarely found in adult guts, this interaction may be the result of *B. apis*-fungal interactions in the diet and where brood are reared. Additionally, the mechanism by which *B. apis* might interact with and/or suppress fungal pathogens is unknown.

Here we examined the potential of *B. apis* to prevent fungal infection in brood and the bacterial genes underlying pathogen defense. To determine the impact of *B. apis* on fungal colonization, we used two different insect pathogens in our assays: *Beauveria bassiana*, a generalist pathogen that infects 70% of insect species, and *A. flavus*, an opportunistic pathogen of honey bee brood. To determine the ability of *B. apis* to inhibit fungal growth *in vitro*, we competed each fungal pathogen with one of five *B. apis* strains, isolated from apiaries in the US (Fig 1a). In the presence of *B. apis* strains, fungal growth was either suppressed or completely inhibited, (Fig 1b). To quantify fungal inhibition, we counted spores of *B. bassiana* or *A. flavus* co-cultured with *B. apis*. The number of spores produced by both *B. bassiana* and *A. flavus*, was reduced by an order of magnitude on average (Fig 1c), showing that *B. apis* can suppress growth of both pathogens.

**Figure 1:**
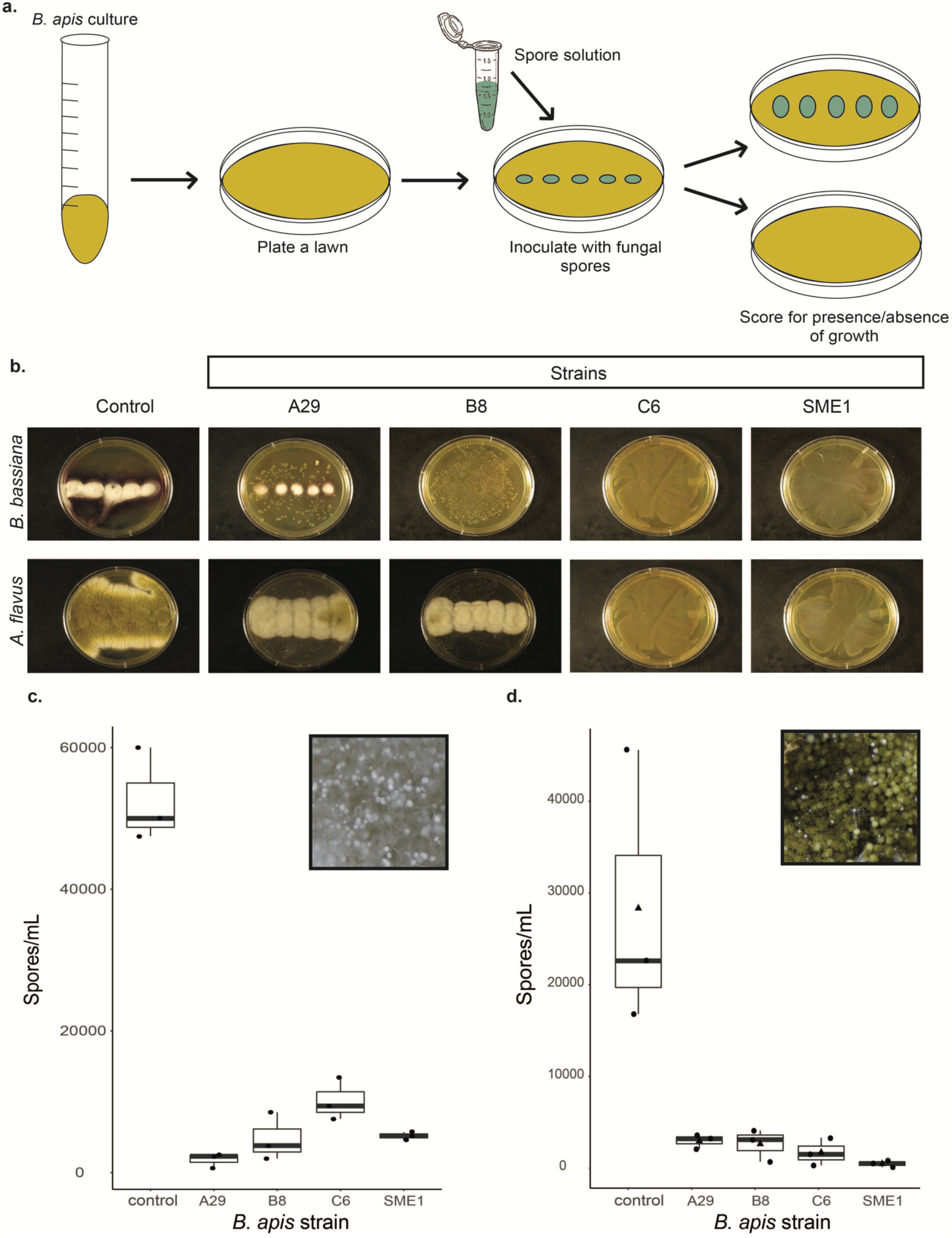
*B. apis* outcompetes fungal pathogens *in vitro.* **a**, The ability of each fungal isolate to grow on a *B. apis* lawns was qualitatively assayed. **b**, Compared to fungal controls, the presence of *B. apis* either suppressed or completely inhibited fungal growth, depending on strain identity. **c**, When co-cultured in liquid media, the presence of *B. apis* strongly reduced the number of spores produced by *B. bassiana* (A29: t = 13.114, df = 2, p = 0.19; B8: t = 11.147, df = 3, p = 0.006; C6: t = 10.121, df = 2.7, p = 0.011; SME1: t = 12.352, df = 2, p = 0.025) and *A. flavus* (A29: t = 2.8807, df = 2, p = 0.40; B8: t = 2.9033, df =2, p = 0.39; C6: t = 3.0137, df = 2, p = 0.37; SME1: t = 3.1679, df = 2, p = 0.34), depending on *B. apis* strain identity.

To test if *B. apis* is capable of preventing fungal infections *in vivo*, we collected larvae from our apiary and reared them on a diet supplemented with either *B. apis* or a sterile media control. Once reared to pupae, the cohort was inoculated with *A. flavus* or a sterile media control and presence of infection was scored until adulthood (Fig 2a). Pupae that were supplemented with *B. apis* as larvae were significantly more likely to resist fungal infection (*χ*^2^ = 14.8, df = 1, p < 0.001), with 66% of the cohort surviving to adulthood with no signs of infection (Fig 2b,c). In sharp contrast, without *B. apis*, no pupae survived to adulthood (Fig 2b, d). Interestingly, in the 34% of *B. apis*-supplemented pupae that succumbed to fungal infection, the number of spores produced was 68% on average (Fig 2e; t = 2.9116, df = 8.4595, p = 0.02). Taken together, these results suggest that the presence of *B. apis* increases the host’s likelihood of survival under fungal challenge, while decreasing the pathogen’s spore load and potential to spread infection to new hosts.

**Figure 2:**
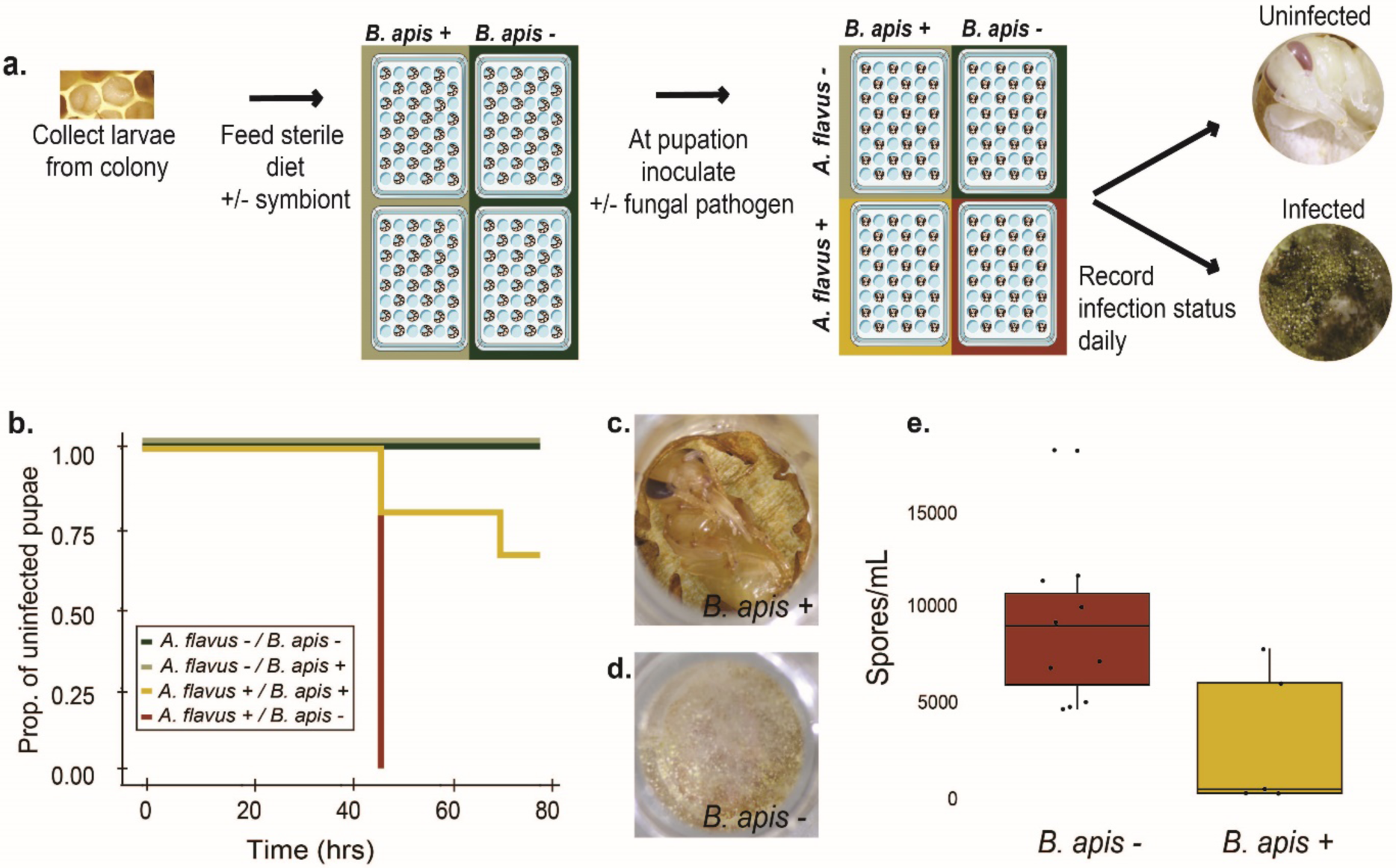
Bee brood supplemented with *B. apis* are less susceptible to infection with *A. flavus.* **a**, First instar larvae (n = 45) collected from the apiary were reared on sterile larval diet +/- *B*. apis (AJP2). Five days after pupation, each pupa was inoculated with 10^3^ spores of *A. flavus* +/- *B. apis* or 0.01% Triton X-100 as a control. **b**, Of the pupae inoculated with *flavus*, those without *B. apis* all showed signs of infection by 48 hrs **d**, whereas 66% of those with *B. apis* never developed infections(*χ*^2^ = 14.8, df = 1, p < 0.001) **c. e**, Pupae with *B. apis* that did become infected had lower intensity infections, producing significantly (t = 5.5052, df = 5.5751, p = 0.002) fewer spores than those without *B. apis*.

To determine if *B. apis* produces antifungal metabolite(s), we incubated fungi in spent media (SM) from *B. apis*, filtered to exclude bacterial cells and normalized for final optical density reached (Fig 3a). Growth of both *B. bassiana* and *A. flavus* were significantly reduced by spent media alone, indicating that *B. apis*-induced changes in the media are sufficient to suppress fungal growth. To eliminate the possibility that fungal inhibition was mediated by acidification of the media, *A. flavus* was cultured in media acidified to pH of 5.0 (the same pH of *B. apis* SM). pH had no significant effect on fungal growth (Fig S3; t = −0.251, df = 35, p = 0.804). Therefore, it is likely that *B. apis* inhibits fungi via secretion of an antifungal secondary metabolite(s). We used antiSMASH^26^ to annotate secondary metabolite gene clusters in the genomes of all *B. apis* strains used in this study and found that all strains have a conserved type 1 polyketide synthase (T1PKS) region. Type 1 polyketide synthases are common among host-associated microbes and produce macrolides which often have antifungal activity ^8,27,28,29^. Additionally, all *B. apis* strains contain an aryl polyene synthesis cluster. The commonly used antifungals amphotericin, nystatin and pimaricin are all polyenes, suggesting that this gene cluster may also contribute to the production of antifungal compound(s). Further functional characterization of these gene clusters will help elucidate whether they play a role in the antifungal phenotype of *B. apis.* Considering the antifungal activity of *B. apis* secreted metabolites *in vitro* and our genomic predictions, it is likely that *B. apis* synthesizes and secretes a metabolite capable of inhibiting fungi.

**Figure 3:**
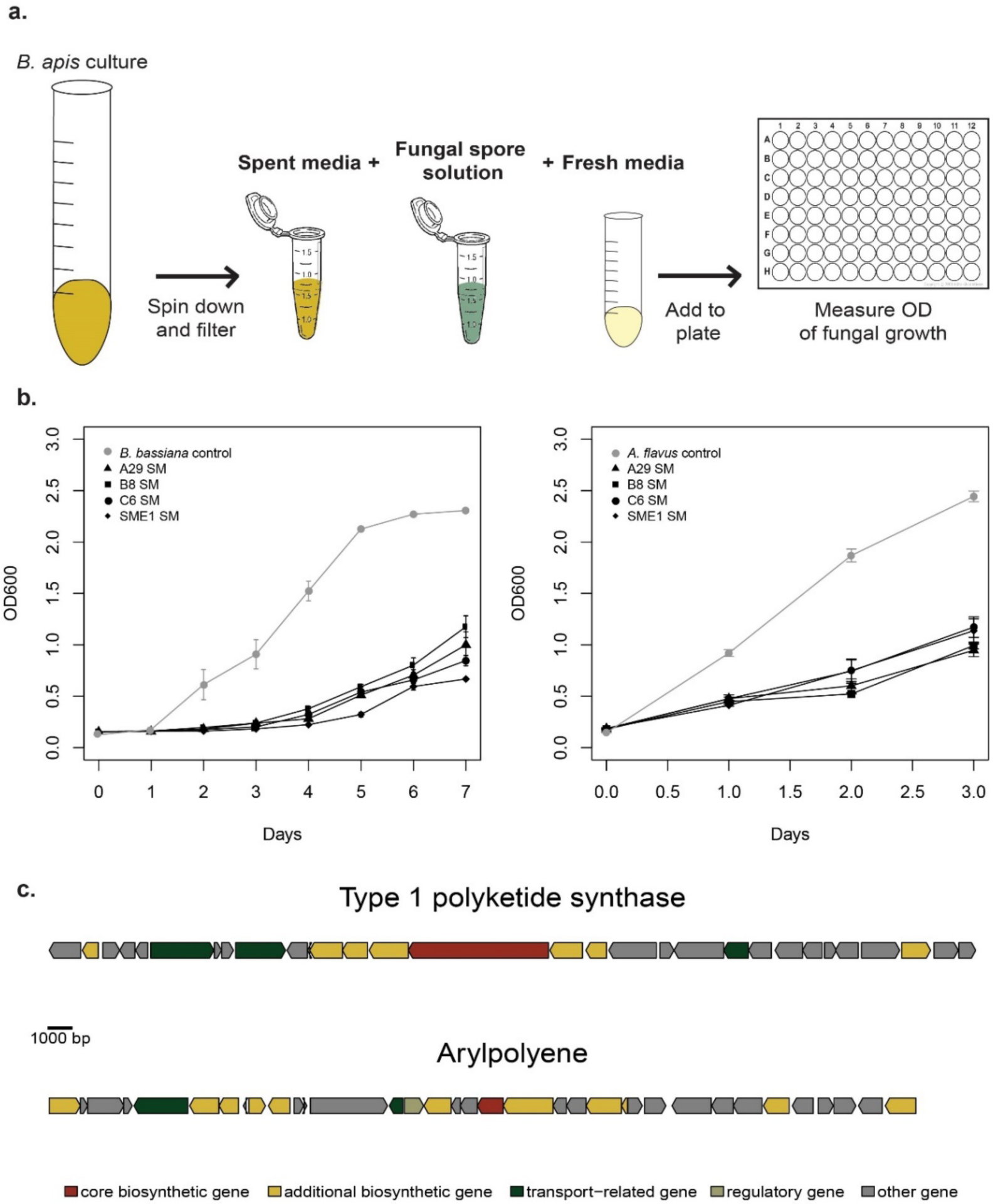
Fungal inhibition is mediated by *B. apis* secreted metabolites. **a**, Spores of fungal isolates were incubated in spent media (SM) from *B. apis* cultures. **b**, The growth of both *B. bassiana* (A29: t = −15.315, df = 119, p < 0.001; B8: t = −13.925, df = 119, p < 0.001; C6: t = −13.202, df = 119, p < 0.001; SME1: t = −11.963, df = 119, p < 0.001) and *A. flavus* (A29: t = −11 .398, df = 59, p < 0.001; B8: t = −13.022, df = 59, p < 0.001; C6: t = −13.282, df = 59, p < 0.001; SME1: t = −11.261, df = 59, p < 0.001) in SM was strongly reduced compared to the control, suggesting secreted metabolites from *B. apis* mediate fungal inhibition. **c**, Genomic architecture of the type 1 polyketide synthase and arylpolyene secondary metabolite gene clusters identified by antiSMASH; gene models are colored based on putative function within the cluster and are oriented to show direction of transcription

Our results provide evidence that a honey bee-associated bacterium, *B. apis*, is capable of suppressing two prevalent insect fungal pathogens both *in vitro* and *in vivo*, likely via the synthesis of an antifungal metabolite. Our *in vitro* results demonstrate antifungal activity in all sampled strains of *B. apis*, with some variation between strains. Analysis of biosynthetic gene clusters present across all strains of *B. apis* revealed two putative regions involved in antifungal production: an aryl polyene synthetase and a T1PKS. Given that a significant proportion of known bacterially-produced antifungals are polyketides^8,27,28,29^, the T1PKS is a promising candidate region.

On the basis of our *in vivo* experiments, supplementing honey bee colonies with *B. apis* may decrease colony losses due to fungal disease. Indeed, in the field, supplementation of *B. apis* is correlated with a reduction in *Nosema* load in adult bees^22^. Beyond decreasing colony losses and fungal load via direct inhibition of fungal infection, the presence of *B. apis* may limit disease transmission by reducing the number of spores produced per infection. In addition, it may suppress adult-specific pathogens, which could be transiently harbored in the larval diet between adult hosts^30^.

Altering the prevalence of pathogenic fungi within managed honey bee colonies could have further ecological consequences. Floral resources shared among diverse pollinators act as transmission centers for fungi, both pathogenic and saprophytic^31^. Species-specific fungal pathogens can be seeded in pollen and nectar sources^32^, after which diverse pollinators, including native bees, can act as vectors to transmit the fungal pathogens to other floral sources, thereby facilitating heterospecific transmission of fungal agents^33^. As a result of reduced spore loads within colonies, the load of fungal pathogens deposited in local floral resources by foragers might also decrease, and perhaps reduce heterospecific transmission and spillover events ^34^.

## Methods Summary

Competition assays were carried out with stationary cultures of *B. apis* normalized to the same OD and 10^3^ spores of either fungal isolate in liquid or solid MRS media. The number of spores produced was counted on a hemocytometer under a light microscope at 40x magnification. Larvae were maintained on UV-sterilized larval diet and supplemented with stationary cultures of *B. apis*. A total of 10^3^ spores of *A. flavus* were added to half the brood, five days into the pupal phase. Presence of fungal infection was scored daily until adulthood. Spent media (SM) of *B. apis* was obtained by spinning down stationary cultures and filtering out remaining bacterial cells using a 0.25 um filter. 10^3^ spores of either fungal isolate were incubated in equal volumes SM and fresh media; OD600 was used as proxy for fungal growth. Genomes for all strains were downloaded from GenBank (see Table 1 for accession numbers) and re-annotated with RAST^35,36^. The resulting GFF files and corresponding genome files were uploaded to antiSMASH ^26^ and results were compared across strains to determine conserved secondary metabolite synthesis clusters.

**Table 1:** Sampling of *B. apis* strains

| species | strain | origin | sample | Genome GenBank accession number |
| --- | --- | --- | --- | --- |
| *B. apis* | AJP2 | NC | nectar | N/A |
| *B. apis* | SME1 | IN | nectar | GCA_009362775.1 |
| *B. apis* | A29 | AZ | larvae | GCA_002917995.1 |
| *B. apis* | B8 | AZ | larvae | GCA_002917945.1 |
| *B. apis* | C6 | AZ | larvae | GCA_002917985.1 |

## Methods

### Isolates and culturing

All bacterial strains of *B. apis* and were obtained by sampling either nectar or larvae (Table 1). Isolates were acquired from our apiary or from Leibniz-Institut DSMZ. All cultures were incubated for 48 hours at 30° C in MRS. Fungal isolates, *B. bassiana* and *A. flavus*, were maintained at 25°C with 80% RH or 34° C with ambient humidity respectively on PDA or MRS agar plates. Spore solutions were prepared by flooding fungal plates with 0.01% Triton X-100, agitating with a cell scraper, and suspending the spores in the solution.

### Competition plates

*B. apis* strains were grown to their maximal OD, and all strains were normalized to the lowest OD value by diluting in fresh media. A lawn of *B. apis* was created by plating 100 µL of normalized culture on MRS agar plates. The plate was then inoculated with 10^3^spores of each fungal isolate and incubated at the appropriate temperature for that isolate. Over the course of three to seven days (depending on isolate) the presence of hyphal/conidia growth was monitored.

### Competition assays

*B. apis* strains were grown to their maximal OD, and all strains were normalized to the lowest OD value by diluting in fresh media. 10^3^ spores of each fungal isolate were incubated in 100 µl of density-normalized *B. apis* culture or 100 µl of fresh media. Fungal growth was monitored daily and once controls showed sporulation, spore counts were quantified for each well via hemocytometer.

### Larval collection and in vivo infections

Late first instars were grafted from our apiary at Indiana University Research and Teaching Preserve into queen cups filled with UV-sterilized worker diet prepared as outlined in Schmel et. al, 2016^37^. *B. apis* supplemented groups were given diet with a ratio of 1:4 stationary (OD=1.0) *B. apis* in MRS to worker diet. This bacterial load was between 2 × 10^6^ and 6 × 10^6^ cells/mL. Control groups were given diet with a ratio of 1:4 axenic MRS media to worker diet. After 5 days in larval diet, pre-pupae were transferred to new wells after either MRS or *B. apis* in MRS was added. Five days into pupal development, individuals were inoculated with 10^3^ spores of *A. flavus* in 0.01% Triton X-100 or an equal volume of 0.01% Trition X-100 as a control. *B. apis*-supplemented groups were co-inoculated with one final dose of the bacterium (10^4^ cells); controls received the same volume of MRS. Presence of infections (as evidenced by hyphae penetrating through the cuticle and/or spore production) was scored daily until adulthood.

### Analysis of biosynthetic gene clusters (BGCs)

Genomes for all strains were downloaded from GenBank (see Table 1 for accession numbers) and re-annotated with RAST^35,36^. The resulting GFF files and corresponding genome files were uploaded to antiSMASH ^26^ and results were compared across strains to determine conserved secondary metabolite synthesis clusters. Gene model figures were visualized and adapted for publication using R^38^.

### *In vitro* antifungal assay

To obtain spent media, strains were grown to their maximal OD (0.6-0.25), and all strains were normalized to the lowest OD value by diluting in fresh media. Cultures were spun down at 9,000 rpm for 5 min and the supernatant filtered through a 0.2 µm filter to remove bacterial cells. Spent media and fresh media were added to a multi-well plate in equal volumes and 10^3^ spores from spore stock solutions were added. Growth was measured daily by assaying *OD*_600_. A positive control included spores in fresh media alone used to compare to treatment groups with spent media. Optical densities of spent media alone were monitored to ensure no bacterial growth occurred. Assay plates were incubated at the appropriate temperature for the fungal isolate used. Since *B. apis* acidifies the media from a pH of 5.5 to 5.0, controls of MRS media reduced to pH 5.0 with HCl were included.

### Statistical analyses

All statistical analyses were performed in R ^38^. Spore counts of fungal isolates in the presence of *B. apis* were compared to controls with unequal variance, two sample t tests; p-values were Bonferroni-corrected for multiple comparisons across strains. *In vivo* infections are displayed as Kaplan-Meier survival curves. *B. apis* +/- infected treatments were compared with a long-rank test using R package, “survminer”^39^. Interactive effects of *B. apis* SM on growth of fungi over time were determined with a generalized linear model of OD, time, and strain identity.

## Data and code availability

All genomic data used in this manuscript are publicly available through NCBI and listed in Table 1.

## Acknowledgements

This work was funded by a *Project Apis m.* grant to ILGN and a USDA NIFA to EAS.

## Author contributions

Conception and design of the work, ILGN and DLM, acquisition, analysis, or interpretation of data, EAS and DLM, drafted and revised the manuscript, DLM, EAS, ILGN.

## Additional information

Supplementary information is available for this paper at: Correspondence and requests for materials should be addressed ILGN.

## Supplemental Data

**Supplementary Figure 1:**
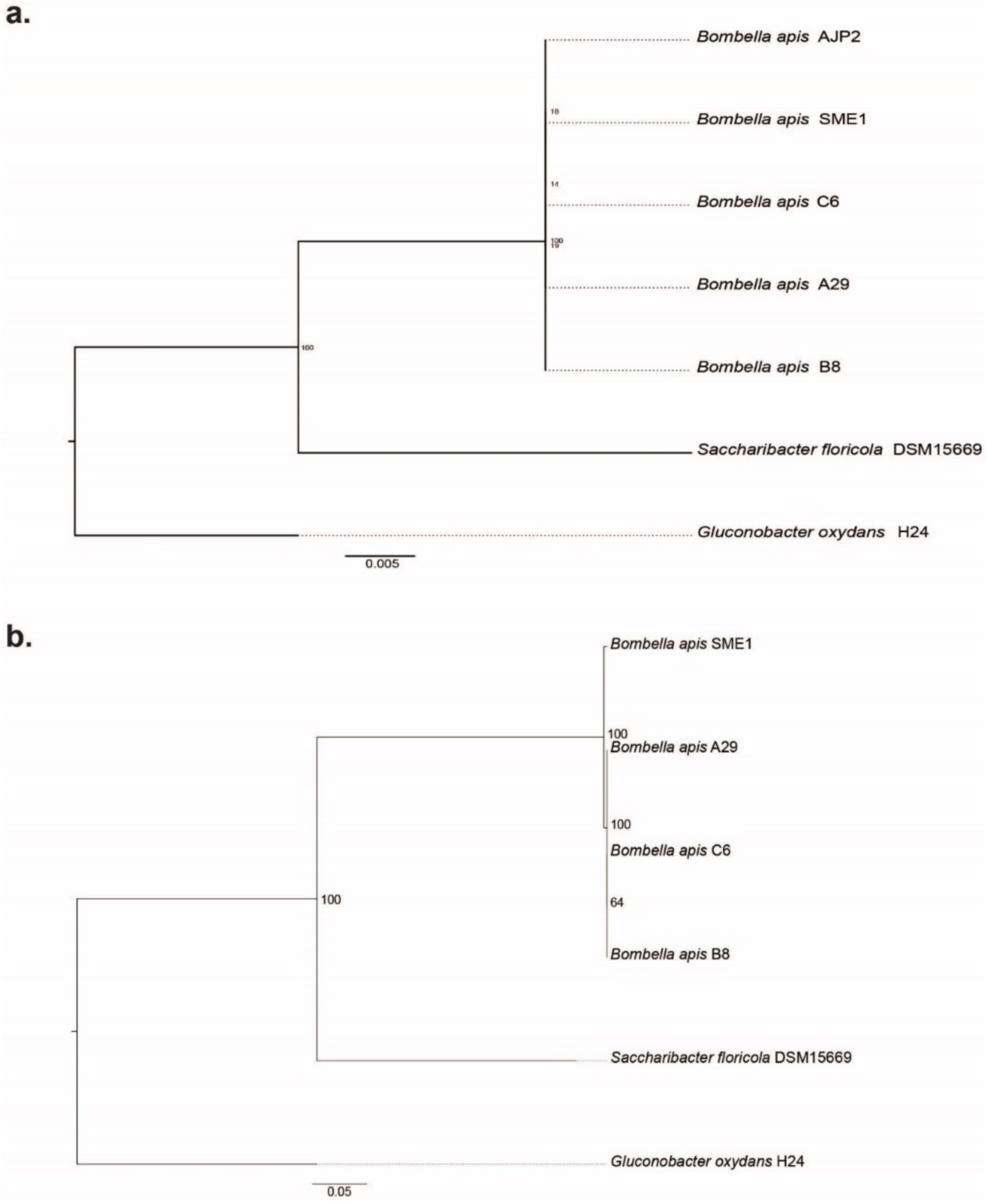
A. Maximum-likelihood 16S rRNA gene sequence tree for strains used in this study. *Saccharibacter floricola* and *Gluconobacter oxydans* were used as outgroups. Sequences were downloaded from GenBank and aligned with the SINA aligner^40^. The tree was constructed with RAxML ^41^ and visualized with FigTree^42^. Numbers at nodes represent bootstrap support from 1000 bootstrap pseudoreplicates. B. Core-ortholog maximum-likelihood phylogeny. All genomes were downloaded from GenBank and core orthologs were identified using OrthoMCL^43^. Alignments of core orthologs were made using MAFFT ^44^and concatenated together. As above, the tree was constructed with RAxML ^41^ and visualized with FigTree ^42^. Numbers at nodes represent bootstrap support from 1000 bootstrap pseudoreplicates.

**Supplementary Figure 2:**
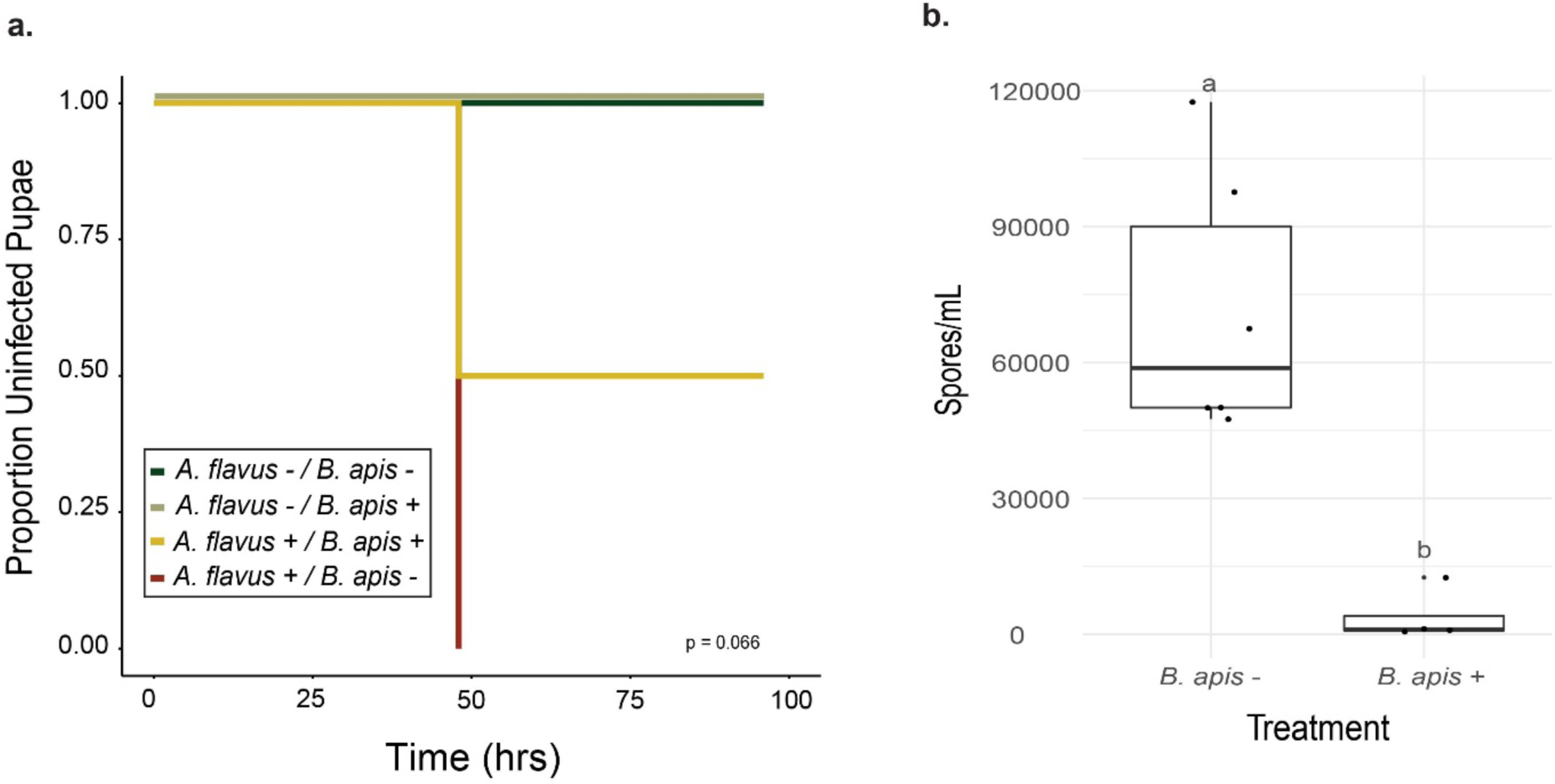
Bee brood are protected from fungal infection, independent of *B. apis* strain identity. **a**, First instar larvae (n = 20) collected from the apiary were reared on sterile larval diet +/- *B*. apis (A29). Five days after pupation, each pupa was inoculated with 10^3^ spores of *A. flavus* +/- *B. apis* or 0.01% Triton X-100 as a control. Pupae supplemented with A29 were more likely to survive to adulthood (*χ*^2^ = 3.4, df = 1, p = 0.07) **b**, Presence of *B. apis* (A29) significantly reduced (t = 5.5052, df = 5.5751, p = 0.001914) sporulation in infected pupae.

**Supplemental Figure 3:**
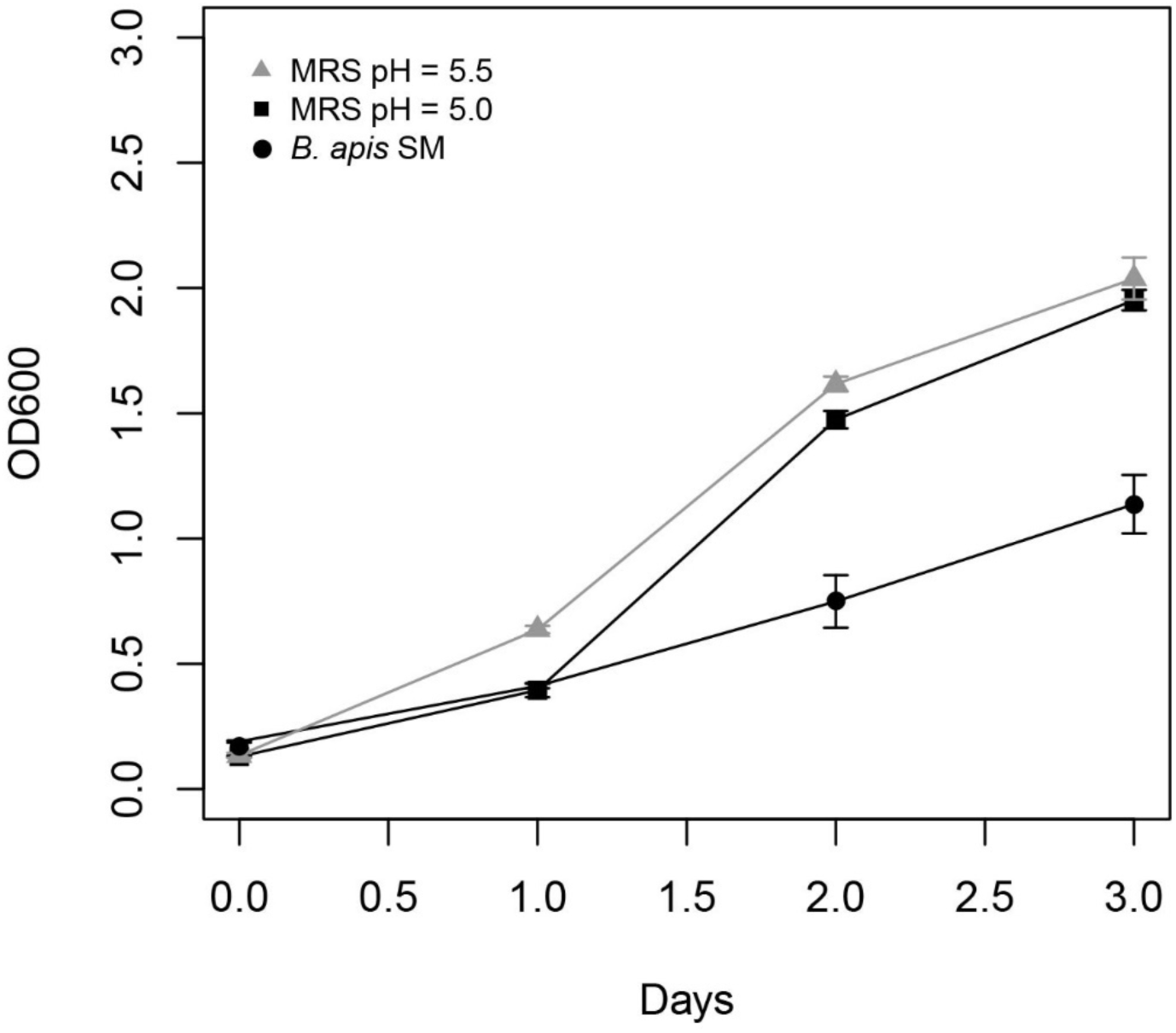
Fungal inhibition by SM is not pH-mediated. *B. apis* (A29) reduces MRS media from a pH of 5.5 to 5.0. Spent media from *B. apis* at pH 5.0 significantly reduced fungal growth (t = −6.111, df = 35, p < 0.001)while MRS media reduced to a pH of 5.0 using HCl did not significantly reduce growth (t = −0.251, df = 35, p = 0.804).

